# NeuroPack: Advanced Learnable Packaging Toolkit Towards Ultra-High Compression Ratio for Volumetric Neuroimaging

**DOI:** 10.64898/2026.10.29.762731

**Authors:** Runkai Zhao, Xiaoxuan Yang, Jiamin Wu, Qihao Zheng, Jiahui Zhang, Jia-Qi Guan, Weidong Cai, Jiu-Lin Du, Lei Bai, Xu-Fei Du, Wanli Ouyang, Guo Lu, Chunfeng Song

## Abstract

High-throughput 3D microscopic imaging generates brain-volume datasets at terabyte to petabyte scale, posing significant storage challenges for large-scale neuroimaging data. In this study, we introduce **NeuroPack**, a hyperprior-driven 3D deep-learning-based packaging toolkit specifically designed for compressing 3D microscopic volumes at the mesoscale. NeuroPack encodes a compact hyperprior stream that conditions its entropy model on local voxel-wise variance, adaptively allocating bits to preserve mesoscale anatomical details for an optimal rate-distortion balance. To ensure reproducible deployment across heterogeneous microscopy datasets and computing environments, NeuroPack implements model integerization for the hyperprior side-information pathway, whose hyper-decoder predicts the Gaussian mean and scale parameters used by the arithmetic coder. By replacing platform-dependent floating-point hyper-decoding with deterministic fixed-point integer convolutions, NeuroPack produces bit-exact entropy models across CPUs, GPUs, local workstations, and cloud servers, thereby preventing decoding drift caused by floating-point rounding, fused-multiply-add ordering, or library-specific kernels. This integerized design preserves rate-distortion behavior while maintaining efficient encoding and decoding for large anisotropic 3D microscopy volumes. Our method enables high-fidelity microscopy reconstruction at 100- to 1000-fold compression across diverse modalities and species. Quantitatively, NeuroPack delivers the strongest overall rate-distortion performance in both BD-PSNR and BD-MSSSIM, with average relative gains of 73.4% and 50.8%, respectively, over the best competing codec. Importantly, NeuroPack preserves image fidelity required for downstream morphological reconstruction and analysis, as validated by simultaneous human annotation of original and decompressed confocal whole-brain zebrafish microscopy images. Together, NeuroPack provides a learning-based compression framework explicitly tailored for microscopy data, establishing a scalable storage solution for data-intensive neuroimaging and AI-empowered biological discovery.

## Introduction

Advanced imaging modalities, including light-sheet microscopy [1][2][3][4][5] and serial scanning electron microscopy [6], produce volumetric datasets that reveal 3D cellular and subcellular structures at unprecedented nanometer to micrometer scales [7] [8][9][10]. However, due to their ultra-high resolution and detailed coverage [11], these imaging datasets frequently reach massive sizes, spanning terabytes for whole-brain mapping and even exceeding petabytes when recording synaptic structures and organelle features [12][13][14]. This substantial data volume imposes considerable challenges in data storage, transfer efficiency, and computational capacity in neuroscientific research [15][16]. Effective volumetric microscopy compression aims to dramatically reduce dataset storage while rigorously preserving biologically essential features, such as neuronal morphology [2][17], precise synaptic connectivity [18][19], organelle integrity [20][21], and cellular boundaries [22][23].

Current conventional image and video compression algorithms can technically be applied to compress volumetric biological data but perform poorly when applied directly to grayscale microscopy volumes. Established on the Joint Photographic Experts Group (JPEG) compression standard, JP3D [24], as part of the JPEG2000 suite [25], has been adapted to process volumetric datasets by incorporating an additional spatial z-axis dimension. This extension leverages 3D discrete wavelet transforms and expands essential concepts such as tiles, precincts, and code blocks into three dimensions, thus facilitating multi-resolution feasibility and enabling region-of-interest access in volumetric datasets [26]. Furthermore, JPEG-XL [27] employing adaptive quantization and variable discrete cosine transform (DCT) represents superior compression for static images. However, when repurposed for 3D microscopy data, the inherent limitations of manually designed transforms become apparent, notably neglecting critical characteristics specific to microscopic imaging, including anisotropic resolutions, intricate structural details, and distinctive redundancy patterns. To effectively address these limitations, specialized compression methodologies explicitly tailored to grayscale volumetric microscopic data are necessary to ensure optimal data fidelity, efficiency, and the preservation of biologically relevant structural information.

An alternative strategy for volumetric data compression designates one spatial dimension as a pseudo-temporal axis and employs conventional video codecs (e.g., H.264 [28] or H.265 [29]). While these codecs can effectively exploit local temporal redundancies through motion estimation algorithms tailored to moving objects in naturalistic settings, they exhibit markable limitations when applied to large-scale 3D microscopic imaging. In particular, standard video compression frameworks operate with a relatively narrow slice-wise receptive field, preventing them from capturing the long-range spatial coherence and structural continuity critical for accurate 3D capture of neuronal arbours. As a result, subtle morphological details spanning adjacent slices may be lost, posing a significant challenge for high-resolution neuroimaging studies where preserving long-context inter-slice features is essential for precise circuit tracing and analysis.

Recent advancements in learning-based image compression models have demonstrated superior performance compared to traditional codecs [30][31][32][33][34]. However, there remains a notable absence of dedicated compression frameworks specifically designed for biological imaging data. Grayscale microscopic volumes often have anisotropic voxel spacing with inhomogeneous intensity [35][36]. Moreover, volumetric data can be extremely large, increasing both the training and inference burden. To handle this disciplinary challenge, in this study, we introduce **NeuroPack**, a deep learning-based neuron volumetric packaging toolkit for high-resolution grayscale microscopic data. By leveraging a data-optimized 3D neural network with entropy modelling, our method exhibits robust generalizability across diverse species and imaging systems, making it a versatile solution for a broad range of neuroimaging data applications. Notably, in preliminary evaluations on BigNeuron [1] and zebrafish private datasets of the light microscope, NeuroPack achieved compression ratios ranging from 10^2^ fold to10^4^ fold, while preserving over high morphological fidelity measured by standard metrics of PSNR and SSIM [37]. This preservation of delicate neuronal structures with low distance discrepancy, even at the whole-brain scale, ensures accurate downstream analyses, from single-neuron annotation to morphological quantification. By significantly reducing the storage footprint, NeuroPack streamlines large-scale data handling, fostering more efficient data exchange and long-term archival and collaborative research. Furthermore, this toolkit supports seamless integration into existing imaging and analysis pipelines and high-throughput compression and decoding processes, enabling the neuroscience community to tailor and refine compression parameters for specific experimental needs. In short, NeuroPack presents an open-source toolkit for the reliable and efficient compression of massive, high-resolution microscopic neuron image datasets.

## Results

### 3D-Hyperprior Supports Versatile Data Compression for Different Species

To evaluate the generalization ability of our proposed 3D-Hyperprior across diverse morphological characteristics, we conduct extensive evaluations on various biological species using Gold166 [1], DIADEM [38], and a proprietary zebrafish dataset (ZebFish-78). These datasets consist of volumetric images captured using optical confocal microscopy. We specifically select training subsets from *Drosophila*, mice, and zebrafish, which focus on representative samples that cover a broad spectrum of morphological variations. To further validate the robustness and generalizability of our approach, we test it on unseen species data, including human and chick samples. This evaluation simulates real-world scientific applications where the compression method must handle unobserved morphological structures without compromising data quality.

Assuming our compression model as a baseline, Bjøntegaard Delta Signal-to-Noise Ratio (BD-SNR) captures the change in signal-to-noise ratio between the compression method and the baseline across the bitrate ranges [39]. A negative value indicates a degradation in performance, while a positive value indicates improvement. Bjøntegaard Delta Multi-Scale Structural Similarity Index (BD-MSSSIM), a perceptual quality metric indicating how well the structural information is preserved across multiple scales [39]. Lower values represent higher degradation. As the baseline, we found that our Hyper3D achieves superior performance across all metrics and species datasets. Traditional compression methods show varying degrees of performance losses, such as JP3D [24], x265 [29], and JPEG-XL [27], with x265 and JPEG-XL often achieving relatively better results, especially in terms of BD-SNR. However, structural fidelity (as reflected in BD-MSSSIM) remains a challenge across all traditional methods, particularly in datasets such as *Drosophila*, zebrafish-ZebFish78 [40], and Chick. JPEG-XL tends to balance signal and structure preservation better than other methods but still falls short compared to our baseline.

Differences in the species and anatomical regions lead to diverse 3D neuron morphologies, which directly result in different image characteristics. To investigate these variations, we conduct a voxel intensity analysis of our test species samples (Figure 1a-b). Voxel intensity distributions vary significantly among species, with distinct patterns that reflect underlying morphological disparities at the microscopic level (Figure 1a). Additionally, the neurite voxels typically exhibit higher intensities compared to neighboring background noise, indicating prominent structural elements. We employ progressive thresholding of voxel intensity values to compute the positive voxels remaining, allowing us to assess the overall structural changes and morphological variations across different species (Figure 1b). Here, *τ* is the voxel-intensity threshold; voxels above *τ* are counted as positive. Therefore, the distinct intensity patterns reflect variations in tissue composition, structural density, and anatomical complexity, which directly impact the efficacy of compression algorithms, which are manifested as observable differences in the 3D volumes, highlighting species-specific features that pose substantial challenges to the development of generalized data compression tools for biological imaging.

**Figure 1.**
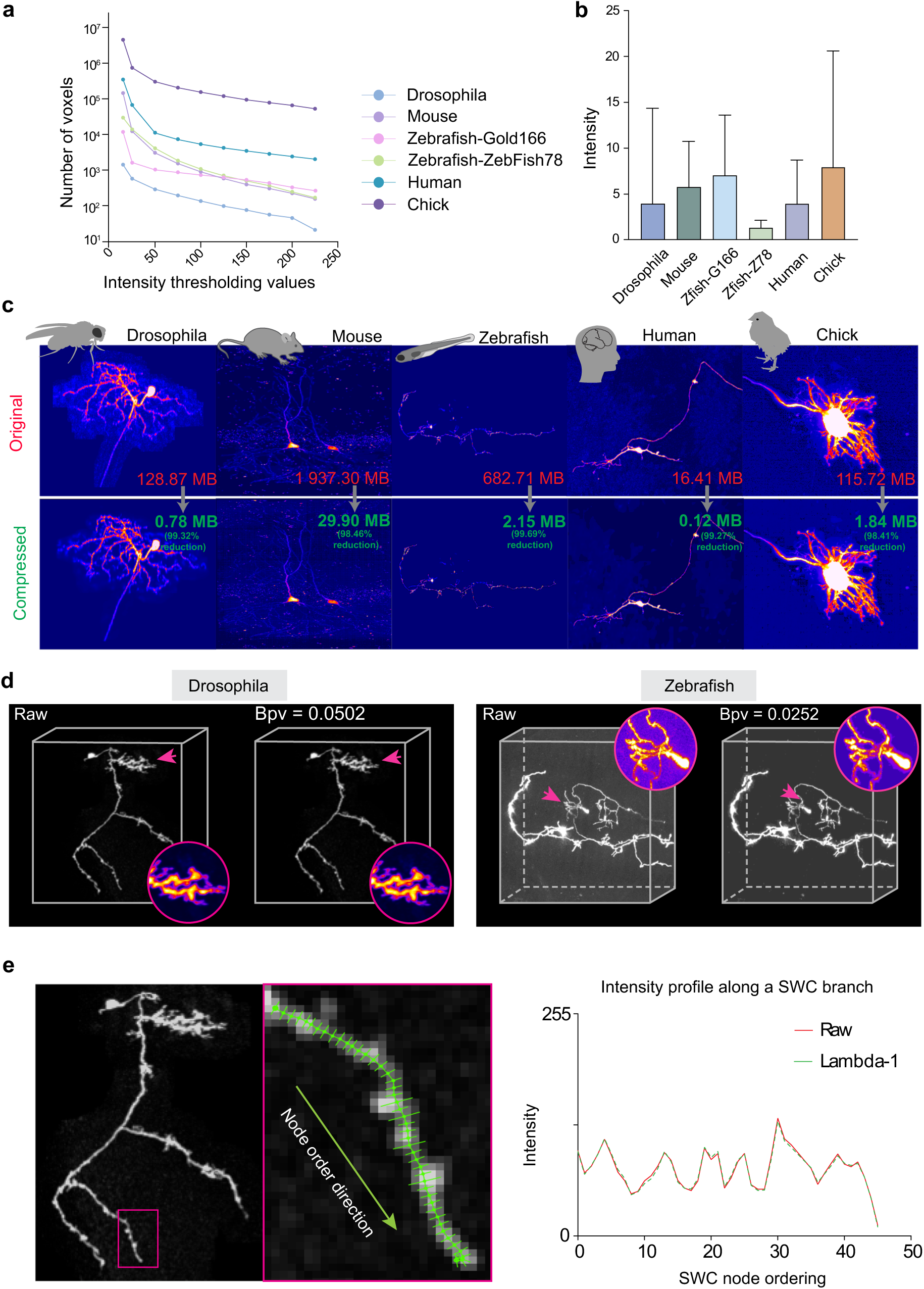
Volumetric light-microscopy compression across species. **a-b,** Inter-species signal characteristics in Gold166 (*Drosophila*, mouse, zebrafish, human, chick) and ZebFish-78. **a,** Voxel-level difference curves: number of voxels whose absolute intensity value exceeds a threshold *τ*, plotted versus *τ*. **b,** Intensity distributions for each species. **c,** Representative volumes per species: original (top) and compressed reconstruction (bottom). Size badges report file sizes for original and compressed data and the achieved bits-per-voxel (bpv). **d,** Zoomed insets from *Drosophila* and zebrafish highlighting preservation of fine morphology (e.g., thin neurites, branch points); annotated regions of interest indicate detailed structures. **e,** Fidelity along a traced neurite: left, volumetric data with an SWC skeleton and a zoomed region; right, the intensity profile along ordered SWC nodes (mean intensity in a local neighborhood), analyzed with the ImageJ SNT plugin [50].

By visualizing in Maximum Intensity Projection (MIP), we compared the compressed microscopic volumes with their corresponding original volumes across various species (Figure 1c). The overall morphological structures of all tested species remain intact and undamaged during the compression process. Additionally, to demonstrate the compression effects on local neuronal structure, we select the test samples of *Drosophila* and zebrafish to visualize the compression results with zoom-in local details. Our method can accurately reconstruct fine image details in close-up visualizations, preserving key structural elements regardless of resolution variations (Figure 1d). This preservation of structural integrity underscores the effectiveness of our compression approach in maintaining critical morphological information across diverse species.

Accurate reconstruction of neuronal morphology relies heavily on maintaining this visual continuity, as any disruptions or inconsistencies in the visual representation can lead to misinterpretation of tracking neurite paths. To evaluate the impact of our compression method on this continuity, we compared voxel intensity changes between the original and compressed volumes along a specific neurite segment (Figure 1e). We use the SWC annotation as a reference and calculate the mean intensity values within node spheres along the neurite. The intensity change trends in the compressed volumes are closely aligned with those in the original volumes, and they have nearly identical intensity values. The compressed volumes generated by our method maintain equivalent visual and morphological quality, which supports reliable neuron reconstruction in both expert and automated analyses.

### 3D-Hyperprior Supports Cross-device Data Compression for 8-bit Microscopic Neuron Volume

Our compression method is designed to handle biological data from various imaging sources. To evaluate the generalization capability of our method, we conducted experiments on microscopic neuron volumes of the mouse brain acquired using the fluorescence Micro-Optical Sectioning Tomography (fMOST) technique [2]. Besides training the model from scratch on fMOST samples, we employed two strategies to leverage pre-trained cross-species knowledge and facilitate adaptation to this new imaging domain. Specifically, we directly applied the pre-trained model to infer the unseen fMOST data. Moreover, we fine-tuned the full model parameters using the fMOST data to further refine the model’s adaptation. These strategies allowed us to evaluate how effectively the model could transfer learned features from previously seen species to unseen new imaging data, highlighting its robustness and versatility in handling biological volumes from different imaging modalities.

We performed a visualization analysis comparing the original dendrite volume with its compressed versions using our fine-tuned model under two different bitrates of 0.0940 and 0.00519 (Figure 2a). A higher bitrate effectively preserves the finer details of the neuronal structures and maintains the overall morphological integrity of the dendrites across these cross-device scenarios. To further evaluate the recovery capability of our method in capturing local morphological details, we examined zoom-in patches of the dendrite volumes. The image with lower bitrates shows signs of quality degradation with noticeable erosions in image clarity and the emergence of discontinuities along the neurite. For quantitative analysis, we compared our compression strategies against x265 and Cheng2020 [31], two methods using volumetric microscopic images (Figure 2b-c). Because the amount of fMOST samples we used here is far less than the dataset we used for training the baseline, training the compression model from scratch cannot effectively capture latent biological patterns and is prone to overfitting. Hence, the models assisted with prior knowledge show better reconstruction accuracy than the model trained from scratch. Across a range of bitrates, direct inference using the pre-trained model delivered performance comparable to other established compression methods, highlighting the strong generalization ability endowed by prior knowledge. The incorporation of this pre-trained knowledge enabled the model to effectively handle diverse biological morphologies without sacrificing fidelity. Furthermore, by employing flexible adaptation techniques, such as fine-tuning, the model could optimally adjust to the new domain data, achieving the highest reconstruction accuracy.

**Figure 2.**
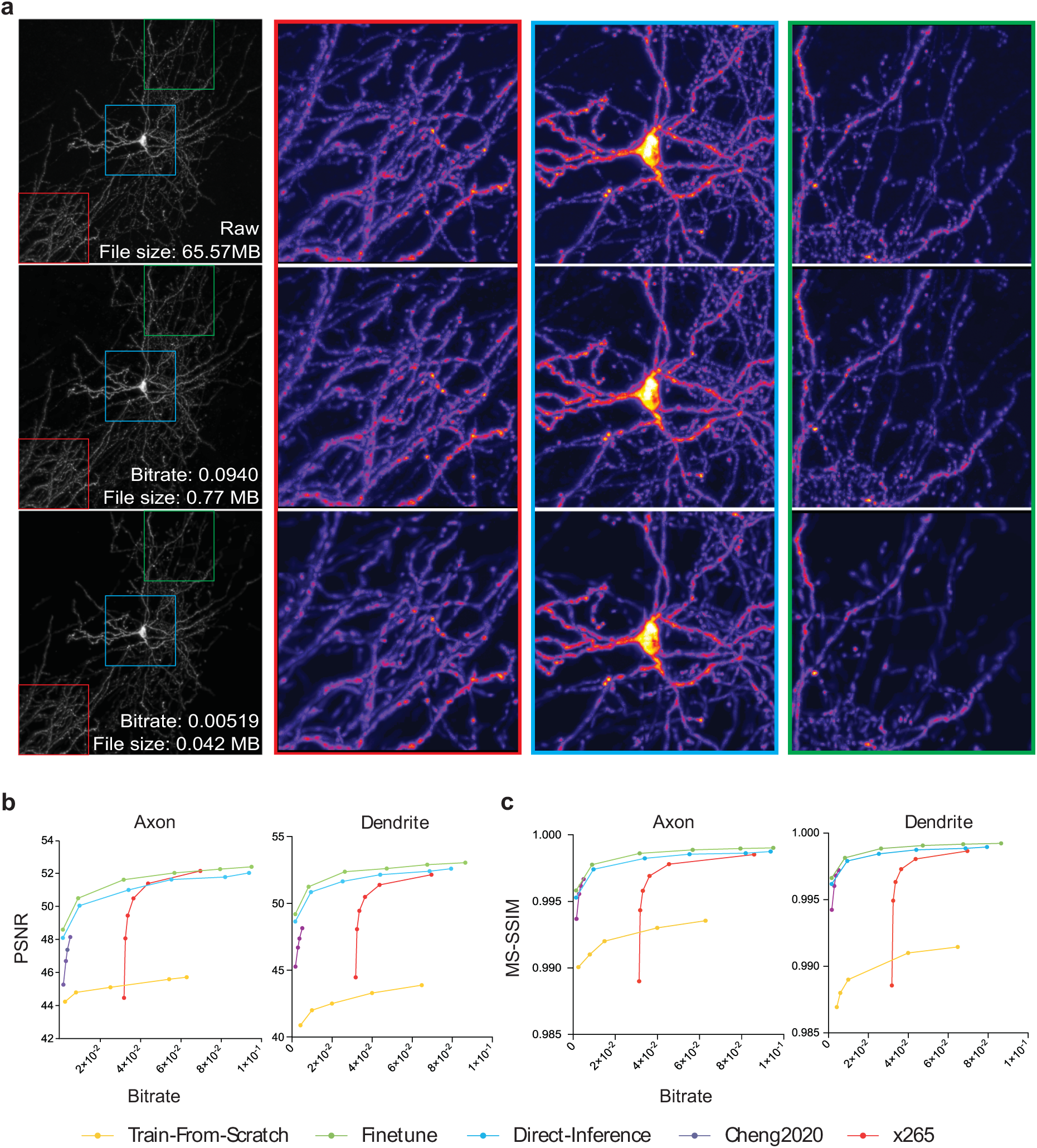
Volumetric light-microscopy compression across acquisition devices. **a,** fMOST volumes shown as maximum-intensity projections at two bitrate points: raw versus compressed reconstructions. Color-coded zoom-in regions of interest (red/blue/green) highlight fine morphological details. **b,** Rate-distortion comparison using PSNR (vertical axis) versus bitrate (horizontal axis, bpv) for a conventional video codec (x265), a learned baseline (Cheng2020), and variants of our method across multiple rates. **c,** The same comparison using MS-SSIM (vertical axis) versus bitrate.

### Affects of NeuroPack on Human Annotation

To assess whether NeuroPack compression compromises biological signal integrity, we employed a two-step labelling procedure. First, we generated baseline labels for each uncompressed image, serving as the ground truth. Next, we held a simultaneous relabeling session, providing both the original and its NeuroPack-compressed counterpart side by side for expert annotation. In Figure 3a, we present visualizations of three annotation variants (Raw, Relabeled, and Compressed) with three standard V3D distance metrics. The color gradient indicates the distance to each soma. These results suggest that overall morphological integrity is preserved under compression. Furthermore, by reducing low-frequency noise, the compression process effectively denoises the images, causing the compressed annotations to more closely resemble the ground truth. In Figure 3b, we evaluate Euclidean distance within weak neurite regions (mean voxel intensity below 20). We find that both relabelled and compressed annotations exhibit a high degree of spatial similarity in these low-intensity areas. Our analysis also indicates that compression does not amplify or exacerbate discontinuities in microscopic data, suggesting it does not adversely affect weaker signal regions. Because bifurcation and terminus nodes are critical for understanding neuronal morphology, we specifically compared distance metrics for these nodes between the relabelled and compressed images against the ground truth. The results reveal that both sets of annotations remain highly similar at these pivotal morphological landmarks. Overall, these findings underscore the effectiveness of NeuroPack in preserving the original neuronal structure with high fidelity.

**Figure 3.**
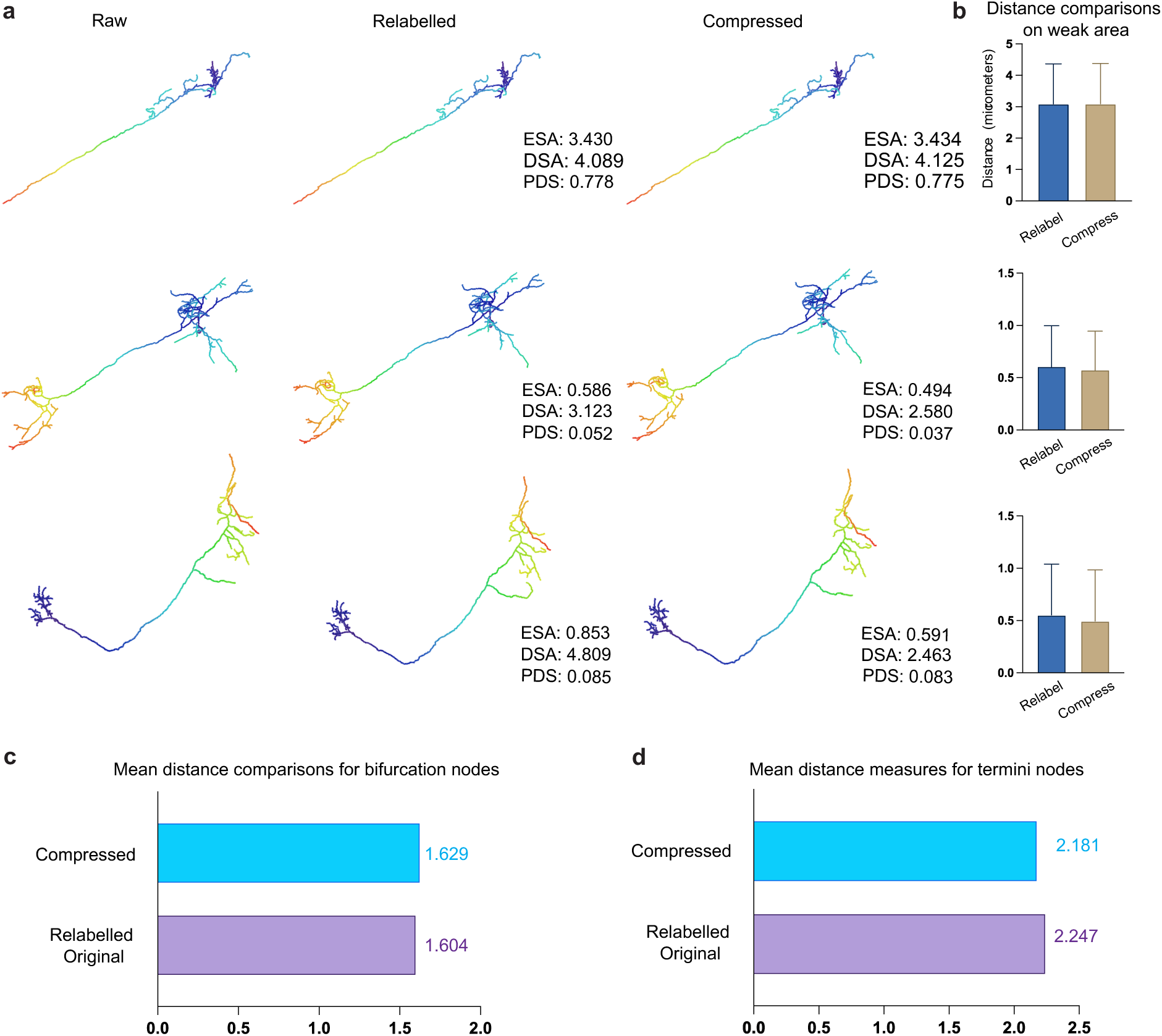
Morphological fidelity of neuronal reconstructions after compression. **a,** Three representative neurons shown as centreline skeletons reconstructed from the raw, expert-relabelled (by using ImageJ annotation tool), and compressed volumes. Colors encode geodesic path length along the tree. Per-cell morphological distance metrics (ESA, DSA, PDS) are reported to the right, showing close agreement between compressed and relabelled references. **b,** Local analyses in weak-signal regions selected by annotators: bar plots compare node-wise distances for relabelled and compressed reconstructions (mean ± s.d.). **c-d,** Dataset-level summaries of node-wise discrepancies at bifurcations (c) and termini (d); mean distances are comparable between compressed and relabelled originals (e.g., bifurcations: 1.629 vs 1.604 voxels; termini: 2.181 vs 2.247 voxels), indicating that compression preserves neurite topology and geometry within annotator variability.

### Cross-device Compression Performance

Cross-platform decoding errors in learned compression of 3D microscopic volumes arise from non-bit-exact floating-point computation in the side-information hyper-decoder ℎ*_s_* , which estimates the Gaussian mean *μ* and variance *σ*^2^ used by the arithmetic coder for the quantized primary latents *ŷ* . Because arithmetic coding is lossless only when encoder and decoder compute identical symbol probabilities, minute platform-dependent discrepancies, stemming from IEEE-754 rounding, fused-multiply-add ordering, kernel implementations, or library versions, can perturb *μ* and *σ*^2^, desynchronize the decoder from the bitstream [41]. This effect could propagate to divergent *ŷ* and corrupted volume reconstructions. To eliminate this nondeterminism, we integerize the trained parameters of ℎ*_s_* and implement all convolutions and transposed convolutions in strictly specified fixed-point/integer arithmetic [42] [43], yielding a bit-exact probability model across CPUs, GPUs, and ASICs and enabling deterministic, portable decoding of large volumetric datasets.

To cross-validate the integerized compression model, we conduct three decoding modes, i.e., integerized on the local host (DII), floating-point on the local host (DIF), and cross-platform (DIC) across anisotropic 3D zebrafish volumes from Zebr300 and ZebrWI datasets [40]. All modes maintained near-lossless structural fidelity at the whole-volume level (MS-SSIM ≥ 0.99 for both datasets), while PSNR largely tracked the intrinsic SNR of each collection (higher for the single-neuron Zebr300; lower for the weak-intensity ZebrWI; Figure 4a). Qualitative assessments corroborated these metrics: thin neurites and branch points were preserved without visible tiling seams or banding artifacts (Figure 4c). Operating at 0.10-0.22 bits per voxel (bpv) yielded substantial storage reductions, for example, a 0.538 Gvox volume achieved ∼75.6× compression at bpv ≈ 0.106, and a 0.813 Gvox volume achieved ∼36.8× at bpv ≈ 0.217, while retaining morphological detail (in-panel gray badges).

**Figure 4.**
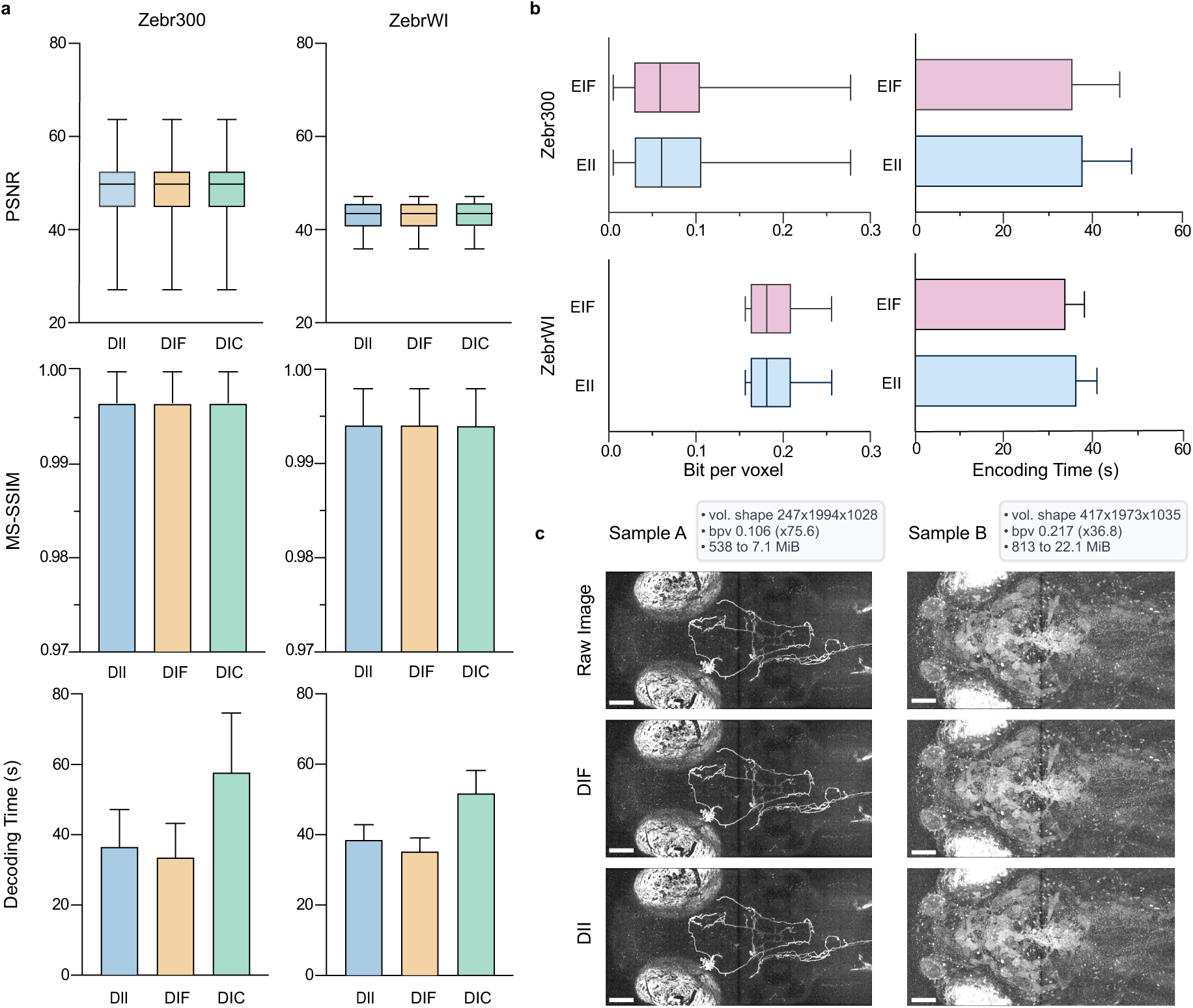
Cross-platform performance evaluations on image fidelity and compression efficiency. **a,** Benchmarking on two zebrafish light microscopy datasets: *Zebr300* (300 volumes of single-neuron staining) and *ZebrWI* (26 volumes of weak-intensity signals). Three decompression modes were compared ---Decoded-in-Integerization (DII), Decoded-in-Floating (DIF), and Decoded-in-Cross-Platform (DIC) ---in terms of image fidelity (PSNR, MS-SSIM) and decoding speed. **b,** Encoding efficiency was assessed by bit-per-voxel and encoding time across two encoding models, EII and EIF. **c,** Representative samples from *Zebr300* (Sample A) and *ZebrWI* (Sample B), showing raw images alongside decompression results. Dataset-specific volume dimensions, compression ratios, and storage sizes are summarized in the gray badges.

On a fixed host, integerized (EII) and floating-point (EIF) encoders produced indistinguishable compression efficiency, because bit-per-voxel (bpv) and reconstruction fidelity were statistically unchanged across modes (Figure 4b). The same held at decode: the computing mode did not affect the achieved compression level (Figure 4a). The primary difference was computational: EII incurred a small wall-clock overhead relative to EIF, reflecting the cost of integerized kernels, while leaving rate–distortion characteristics unaffected.

Notably, a single bitstream is decoded by both DII and DIF, ensuring numerical reproducibility on integer paths (important for auditability and multi-site replication) while allowing floating-point decoders to integrate seamlessly with GPU-native pipelines. Collectively, these findings indicate that sub-0.25 bpv operating points are sufficient to preserve micron-scale structure in large, anisotropic light-microscopy volumes, with DII/DIF and EII/EIF covering complementary corners of the fidelity–throughput–portability trade space for scalable neuroimaging workflows.

## Discussion

In summary, we have developed a 3D-Hyperprior compression strategy, which we refer to as NeuroPack when applied to human annotation data, to efficiently preserve the morphological fidelity of neuron volumes across multiple species and imaging modalities. By leveraging deep learning-based latent representations, our approach reduces low-frequency noise and maintains critical structural details, thereby improving downstream tasks such as neuron reconstruction and expert annotation. Through extensive evaluations on *Drosophila*, mouse, zebrafish, human, and chick datasets, we have demonstrated that morphological landmarks (e.g., bifurcation and terminus nodes) remain intact, even in low-intensity regions.

Our method is versatile and accommodates further enhancements. For instance, it can readily incorporate new pre-trained models trained on additional species or imaging techniques, thereby further improving its generalization to previously unseen data. In practice, compression performance may be affected by factors such as the inherent quality of raw microscopic images, tissue composition, and labelling sparsity. More robust training on diverse data, as well as the introduction of advanced noise reduction or domain-adaptation algorithms, can further mitigate these challenges.

Moreover, although our current approach preserves morphological integrity effectively, compression artifacts may still appear in extremely low-bitrate scenarios or with highly variable imaging conditions. In these cases, local discontinuities might emerge and warrant careful scrutiny by domain experts. Fine-tuning or adapting the model to these specific imaging conditions can help reduce potential structural inconsistencies and improve reconstruction quality.

Additionally, advanced strategies to distinguish complex branching patterns can be introduced, thereby extending NeuroPack’s applicability to more intricate neuronal circuits. For datasets with limited imaging quality, the compression’s denoising benefits could be a double-edged sword; while background noise is effectively reduced, detail in ambiguous regions might also be smoothed out. Consequently, imaging and labelling improvements remain critical for maximizing reconstruction accuracy across large-scale neuron datasets.

Finally, the ability to combine NeuroPack with other specialized neuron reconstruction pipelines, particularly those that incorporate multi-round proofreading or crowd-based annotation, can further accelerate the throughput of large-scale neuron mapping. By preserving key morphological features, our method reduces the potential need for corrective edits, streamlines the expert proofreading process, and fosters more reliable reconstructions. As a result, NeuroPack stands as a robust framework for neuron data compression, with significant potential to enhance next-generation neuroscientific studies involving diverse biological species and heterogeneous imaging techniques.

## Methods

### Models of Comparison

**JP3D** is an encoder designed for the compression of 3D data, utilizing a 3D wavelet transform for efficient data compression. It allows for the direct input of 3D image data, although there are limitations on data size. When the dimensions of a single image are too large, attempting to input the entire image may result in memory errors; therefore, we divide the image into patches of size [128,128, 64] for input. JP3D supports direct adjustment of the compression ratio, which can be set within the range of 1 to 3000, and this ratio does not self-adjust based on the compressed data.

**x265** is an open-source and free HEVC (High Efficiency Video Coding) video encoder. It can encode 3D image data by treating one dimension as the temporal dimension, with each slice considered as a video frame. The video encoder includes features such as intra-frame prediction, inter-frame prediction, transformation, quantization, and entropy coding. x265 supports video format input. We convert the 3D images losslessly into YUV video format and then divide them into patches of size [128,128, 64] for compression using x265. We selected the "fast" encoding mode and adjusted the CRF (Constant Rate Factor) parameter to achieve the desired changes in bitrate.

**JPEG XL** is a practical approach focused on efficient compression of high-quality images, representing the optimal choice among traditional image encoders. Since JPEG XL only supports 2D image encoding, we must split the 3D image into multiple 2D images before inputting them into the encoder. We perform the splitting along the z-dimension of the 3D image, treating every three slices as a single-color image for separate cutting and encoding. We adjust the quality parameter q to achieve the desired changes in bitrate.

**cheng2020-anchor** is a learning-based 2D image codec that employs a hyperprior and a context model for joint encoding. The transform module in our model is consistent with that of cheng2020-anchor. Since cheng2020-anchor only supports 2D image input, we split the 3D image along the z-channel to obtain color image slices. The pre-trained model and test code for cheng2020-anchor are sourced from the open-source library compressAI. We utilize the corresponding pre-trained checkpoints for various target bitrates.

**mbt2018-mean** is a learning-based 2D image codec that employs hyperprior encoding, which serves as the fundamental architecture for learning-based image compression. The underlying structure of our model is consistent with that of mbt2018-mean. Since mbt2018-mean only supports 2D image input, we also split the 3D image along the z-channel to obtain color image slices. The pre-trained model and test code for mbt2018-mean are sourced from the open-source library compressAI. We utilize the corresponding pre-trained checkpoints for various target bitrates.

### NeuroPack Architecture Details

We extend the canonical Hyperprior-2D codec [44], [45] to anisotropic 3D microscopic volumes with a mean-scale Gaussian hyperprior as shown in Figure 5. We target grayscale microscopy volumes in which all axes are visually important with 8-bit intensities (0-255). The main analysis transform *g_a_* encodes a single-channel input *x* ∈ *R*^1×*D*×*H*×W^ into a latent *y* ∈ *R*^128×*D*/8×*H*/16×W/16^, where ℝ is the real-valued tensor space; D, H, and W denote depth, height, and width; and y is the 128-channel primary latent. The transform uses stacked 3D residual convolutional blocks ( 3 × 3 × 3 ), GDN3D nonlinearities [46], and strided downsampling. To accommodate lower axial resolution typical of biological acquisitions, one block applies anisotropic stride of 1 × 2 × 2, and overall the pyramid reduces resolution by × 16 × 16 (depth×height×width) via strides (2, 2, 2), (2, 2, 2), (1, 2, 2), and a final (2, 2, 2) stage. Skip connections use 1 × 1 × projections. The synthesis transform *g_s_* mirrors this design with residual blocks, transposed 3D convolutions (including an anisotropic 1 × 2 × 2 upsampling stage), and IGDN3D [46] to produce the reconstructed volume *x̂*.

**Figure 5.**
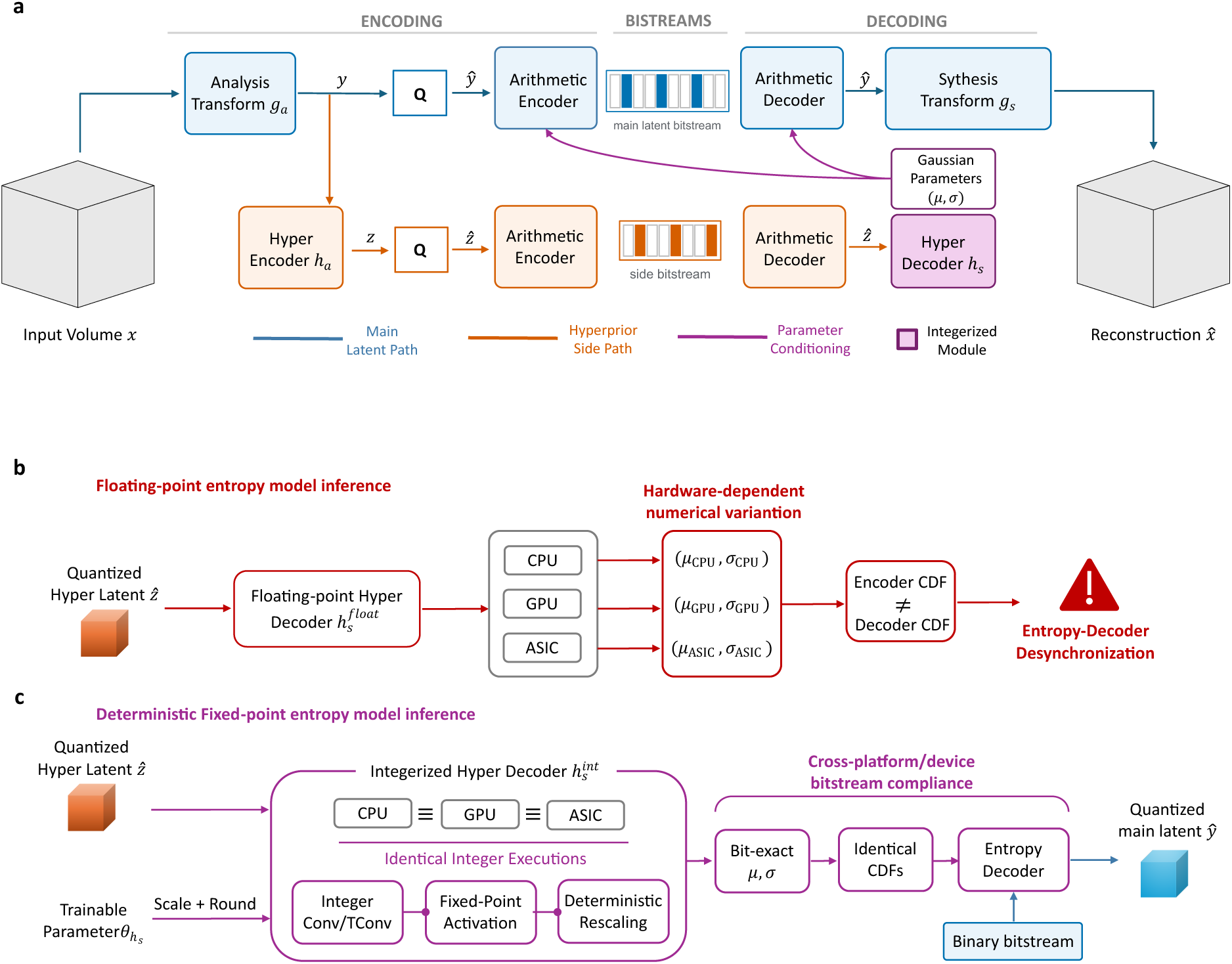
NeuroPack architecture and deterministic cross-platform entropy decoding. **a,** The input volume *x* is mapped by the analysis transform *g_a_* to primary latents *y*, which are quantized and entropy-coded into the main bitstream. The hyper-analysis transform ℎ*_a_* maps *y* to hyperlatents *z*, which are transmitted through a smaller side bitstream. The decoded hyper latents *ẑ* are processed by the integerized hyper-synthesis transform 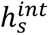 to generate spatially varying Gaussian mean *μ* and scale *σ*, which condition the main-latent entropy encoder and decoder. The decoded latents *ŷ* are transformed by *g_s_* to reconstruct *x̂*. **b,** Floating-point evaluation of ℎ*_s_* can produce hardware-dependent numerical variations that alter entropy parameters, scale indexes or symbol offsets, resulting in a decoder-side probability model that differs from that used during encoding and thereby risking entropy-decoder desynchronization. **c,** In the fixed-point pathway, the trained parameters and intermediate features of ℎ*_s_* are quantized, and all convolution, transposed-convolution, activation and rescaling operations follow specified integer-arithmetic rules. This produces bit-exact entropy parameters and identical discrete cumulative distribution functions across CPU, GPU and ASIC implementations, enabling deterministic decoding of the same main bitstream. Blue, orange and magenta denote the main-latent pathway, hyperprior side pathway and entropy-parameter conditioning, respectively.

The hyperprior pathway further models local uncertainty in the primary latents *y*. The hyper-encoder ℎ*_a_* further encodes *y* into the compact hyperlatent *z* ∈ *R^C^*^×*D*/32×*H*/64×W/64^, where C is the channel count; it is entropy-coded through a bottleneck. The hyper-decoder ℎ*_s_* maps the quantized hyperlatent *ẑ* into spatially varying mean and scale fields (*μ*, *σ*) for each channel of the quantized latent *ŷ*; μ is the mean and σ the positive scale (standard deviation), yielding locally adaptive Gaussian probability models for arithmetic coding. Concretely, ℎ*_s_* emits paired mean/scale feature maps. We employ LeakyReLU activations in selected blocks and use a non-negativity reparameterization for σ to stabilize both normalization and scale estimation.

### Model Integerization

We integerize the hyper-decoder ℎ*_s_* because it directly generates the per-voxel Gaussian parameters that drive the arithmetic (range) coder for *ŷ* . These must be bit-exact on both encoder and decoder. Even subtle floating-point differences alter symbol probabilities and desynchronize decoding. Making ℎ*_s_* fixed-point guarantees identical (*μ*, *σ*) across CPUs/GPUs/ASICs, ensuring bitstream compliance.

Concretely, the fixed-point path uses input-feature scale *K*_1_ and parameter scale *K*_2_. Here, *x*, *w*, and *b* denote input features, weights, and biases with subscripts *f* and *i* indicate floating-point and integer forms. The integer forms follow Equations (1)-(3):

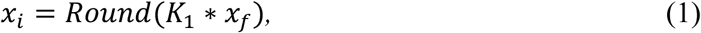

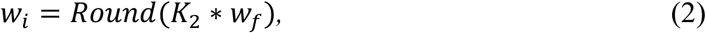

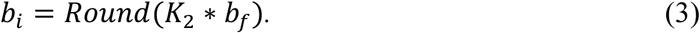

Equation (4) applies integer convolution or transposed convolution, where *Conv* is the operator and *y_i_* the output at scale *K*_1_:

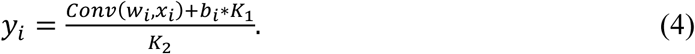

At the interface, outputs are rescaled (and, when needed, rounded) to recover *ẑ* or the Gaussian parameters supplied to the entropy model. This three-step procedure consists of (1) input quantization, (2) weight and bias quantization, and (3) integer convolution, yielding a bit-exact realization of ℎ*_s_* without altering rate-distortion behavior, thereby eliminating platform-dependent decoding errors.

### Training Configurations

We train the 3D hyperprior codec with a standard rate–distortion objective on randomly sampled 3D patches. Volumes from the target light-optic microscopic collections are read as 8-bit TIFFs, intensity-normalized to [0,1], and cropped into 64 × 128 × 128 patches. All-zero patches are rejected to avoid empty inputs. Training is purely self-reconstruction, comparing the reconstructed output *x̂* with the original input *x*. The training loss is formulated as a weighted sum of bit-per-voxel loss and MSE loss with *λ* = 0.1 as the rate-distortion weighting coefficient. Optimization procedure uses Adam with an initial learning rate of 1 × 10^−4^, and a StepLR schedule decays the learning rate by 10 × at 0.75 of the total training epochs [47].

## Data Availability

The exemplar zebrafish volumetric microscopic data will be published via Zenodo.

## Code Availability

All code and sample zebrafish volumes used in this study is available at https://github.com/SciDataOcean/NeuroPack. We mainly implemented CompressAI Torch library [48] as main code framework for modelling building and experimental conduct. We used Vaa3D [49] and ImageJ SNT [50] software for visualization and analysis.

**Table 1.** BD-PSNR Compression Performance of NeuroPack compared to state-of-the-art codecs. Bjøntegaard Delta peak signal-to-noise ratio (BD-PSNR, unit: dB) [51] is reported relative to JP3D, which is used as the reference codec. Higher values indicate better rate–distortion performance. The best results for each dataset are highlighted in **bold**, and the second-best results are <u>underlined</u>.

|  | *Drosophila* | *Mouse* | *Zebrafish*<br>(*Gold166*) | *Zebrafish*<br>(*ZebFish78*) | *Human* | *Chick* |
| --- | --- | --- | --- | --- | --- | --- |
| JP3D [24] (**baseline**) | 0 | 0 | 0 | 0 | 0 | 0 |
| x265 [29] | 7.240 | -0.904 | **3.877** | <u>8.939</u> | 0.546 | <u>2.365</u> |
| Cheng20 [31] | 3.222 | <u>-0.188</u> | -1.148 | 5.930 | <u>0.657</u> | 1.234 |
| Hyper2D [45] | 4.529 | -0.494 | -3.156 | 4.833 | 0.510 | 0.529 |
| JPEG-XL [27] | <u>7.505</u> | -0.897 | 0.960 | 6.838 | 0.218 | 1.209 |
| NeuroPack (**Ours**) | **14.335** | **3.101** | <u>3.339</u> | **11.772** | **2.581** | **3.134** |

**Table 2.** BD-MSSSIM compression performance of NeuroPack compared with state-of-the-art codecs. Bjøntegaard Delta multi-scale structural similarity (BD-MSSSIM, unitless) is reported relative to JP3D, which serves as the reference codec for most datasets. For Mouse and ZebFish78, JP3D results were excluded due to prohibitive runtime; in these cases, x265 and Hyper2D, *both with relatively lower performance*, were used as alternative references. Higher BD-MSSSIM values indicate better perceptual rate–distortion performance. The best result for each dataset is shown in **bold**, and the second-best result is <u>underlined</u>.

|  | *Drosophila* | *Mouse* | *Zebrafish (Gold166)* | *Zebrafish (ZebFish78)* | *Human* | *Chick* |
| --- | --- | --- | --- | --- | --- | --- |
| JP3D [24] | 1.197 | N/A | 1.945 | N/A | -1.353 | -9.254 |
| x265 [29] | 6.974 | 0.551 | **7.332** | <u>7.500</u> | -0.378 | 1.426 |
| Cheng20 [31] | -0.404 | <u>0.679</u> | 2.810 | 1.014 | <u>0.239</u> | 0.529 |
| Hyper2D [45] (**baseline**) | 0 | 0 | 0 | 0 | 0 | 0 |
| JPEG-XL [27] | <u>9.538</u> | 0.007 | <u>6.671</u> | 6.919 | -0.097 | <u>1.733</u> |
| NeuroPack (**Ours**) | **15.084** | **0.801** | 6.480 | **10.074** | **2.669** | **2.254** |

## Acknowledge

This work was supported by the Shanghai Artificial Intelligence Laboratory, and the Lingang Laboratory (Grant Nos. LGL-1630-02 and LGL-1987-19); the National Natural Science Foundation of China (Grant Nos. 62320106010 and 32321003); and the Brain Science and Brain-like Intelligence Technology -National Science and Technology Major Project (Grant Nos. 2021ZD0204500 and 2021ZD0204502). Additionally, we are grateful to Researcher Wenzhe Zhong (Institute of Neuroscience; State Key Laboratory of Neuroscience; Center for Excellence in Brain Science and Intelligence Technology, Chinese Academy of Sciences) for preparing and annotating the microscopy data.

## Notes

### Competing Interest Statement

The authors have declared no competing interest.

